# Habitat use differences mediate population threat exposure in white sturgeon

**DOI:** 10.1101/2022.08.31.505999

**Authors:** Jonathan A. Walter, Gabriel P. Singer, Daniel C. Reuman, Scott F. Colborne, Lawrence W. Sheppard, Daniel R. O’Donnell, Nat Coombs, Erin E. Tracy, Myfanwy Johnston, Emily A. Miller, Anna E. Steel, John T. Kelly, Nann A. Fangue, Andrew L. Rypel

**Author notes:** Corresponding Author: Jonathan A. Walter Center for Watershed Sciences University of California, Davis 1 Shields Ave Davis, CA 95616.

## Abstract

Understanding intraspecific variation in habitat use, particularly of long-lived fishes across multiple life history stages, is core to improved conservation management. Here, we present results from a synthesis of acoustic telemetry data for large juvenile and adult white sturgeon (*Acipenser transmontanus*) from 2010 to 2017 in the San Francisco Estuary and Sacramento River ecosystems. We focused primarily on uncovering spatial patterns of inferred habitat occupancy across life stages, and on linking habitat use to population threats. We found substantial differences in habitat use across individuals and over time that was related to fish age class. However, differences in habitat use were not explained by fish sex or water year flow conditions. We estimated an index of angling exposure, which showed that fish entering reproductive maturity, which historically were of harvestable size, were detected less often than other sizes in areas with high angler pressure, suggesting possible behavioral avoidance of areas of high angler pressure. Additionally, we used historical data to evaluate potential exposure of white sturgeon to a severe red tide event in late summer 2022. We found that >50% of reproductive-age fish may have been residing in areas affected by the red tide. Future monitoring and management of white sturgeon might benefit from examining multiple phases of white sturgeon life history. For example, additional tracking studies could improve understanding of juvenile habitat use, adult survival rates, patterns of anadromy, and cross-basin habitat utilization.

## Introduction

Understanding how patterns of fish habitat use vary within and among populations is essential to conservation and management (Crossin et al. 2017). In fishes, habitat use and movement may differ among individuals of the same population according to life stage, sex, and behavioral types (Hutchings and Gerber 2002; Moyle 2002; Bade et al. 2019; Colborne et al. 2019). Anadromy is an obvious example of habitat use and movement differing across life stage for the same individuals. As an example of sex-related differences in movement behavior, in a population of brook trout (*Salvelinus fontinalis*) on Cape Race, Newfoundland, Canada, male fish were more likely to disperse and tended to move more than twice as far as females (Hutchings and Gerber 2002). In other examples, distinct movement behavior may be exhibited by groups that do not differ according to characteristics such as age or sex. In the Laurentian Great Lakes, potamodromous and adfluvial lake sturgeon (*Acipenser fulvescens*) exhibited within-population differentiation of movement patterns, with distinct groups occurring in different areas but not differentiated by sex or age (Kessel et al. 2018; Colborne et al. 2019). These behavioral patterns can reflect adaptive functions such as exploiting resources, securing mates, and minimizing competition (Chapman et al., 2012). Differences in habitat use and movement, such as partial migration, may also stabilize populations through a bet-hedging mechanism against local catastrophe, often referred to as the portfolio effect (Secor 1999; Chapman et al. 2012; Gomez et al. 2015; Kessel et al. 2018).

Acoustic telemetry is a widely used technology for inferring habitat use and movements of fishes with diverse lifecycles (Donaldson et al. 2014; Hussey et al. 2015). Although telemetry studies have long provided insights for fish ecology and conservation (Crossin et al. 2017), more recent developments such as passive receiver array design, improvements in transmitter size and longevity, and establishment of cooperative networks of investigators and government agencies (e.g., GLATOS, the Great Lakes Acoustic Telemetry Observation System, Krueger et al., 2018; OTN, the Ocean Tracking Network, Iverson et al., 2019) enable large-scale, long-term studies across many individuals and across investigating institutions. Leveraging emerging technologies for better long-term tracking of individuals is particularly important for highly mobile and long-lived fishes like sturgeon (Altenritter et al. 2018; Colborne et al. 2019, 2022; Breece et al. 2021).

White sturgeon (*Acipenser transmontanus*) is a long-lived, semi-anadromous fish species inhabiting large river basins along the Pacific coast of North America (Moyle 2002). White sturgeon have for several years been a California state ‘Species of Special Concern’ due to anthropogenic threats including extensive habitat degradation and loss, notably through channel modification, dams, and water diversions (Moyle 2002). A recent analysis estimated that the population growth rate of white sturgeon in the San Francisco Estuary and Sacramento and San Joaquin river systems may be below replacement levels and, consequently, the population may be in decline (Blackburn et al. 2019). In July, 2024, white sturgeon became a candidate for listing under the California Endangered Species Act (CESA). This potential action was spurred in part by an unprecedentedly severe and widespread red tide event (also known as a harmful algal bloom or HAB) in the San Francisco Estuary that occurred during August 2022. This event produced a fish kill, the scope of which was difficult to quantify. However, significant mortality of white sturgeon occurred; at least 864 sturgeon (green, white, and unidentifiable) carcasses were reported, and this is suspected to be only a fraction of total mortality due to the red tide (Calfornia Department of Fish and Wildlife 2023).

In addition to the impacts of environmental changes and disturbances, white sturgeon have historically supported a significant recreational and harvest-oriented fishery in California, although white sturgeon harvest is currently illegal under emergency regulations while it is evaluated for listing under the CESA. From 2007 through 2013, the legal harvest slot limit was 117–168 cm Total Length. In 2013 and for the remainer of the study period, slot limit measurements were change to Fork Length and the upper and lower limits adjusted to 102–152 cm to reflect fish of equivalent size to the prior limit. Like other species that take several years to reach sexual maturity (Reynolds et al. 2005; Pinsky et al. 2011), white sturgeon are vulnerable to overfishing (Blackburn et al. 2019; McLean et al. 2020). Despite consistent monitoring of the fishery, habitat use and movements of white sturgeon in the San Francisco Estuary and Sacramento River system are relatively little studied (but see, e.g., Johnston et al., 2020; Miller et al., 2020), leaving open questions about their ecology, patterns of habitat use, and effective approaches to improving habitat and fishing conditions to reverse declining abundance trends.

This study synthesized seven water years of acoustic telemetry records to study patterns of habitat use by large juvenile and adult white sturgeon in the San Francisco Estuary and Sacramento River system. We first asked (1) whether white sturgeon individuals exhibited differentiable patterns of habitat use. Finding evidence that white sturgeon differed in habitat use patterns, we next asked (2) whether movement and habitat use patterns were related to fish age class, fish sex, and hydrologic conditions. To examine conservation and management implications for these differences, we also asked (3) whether groups of white sturgeon displaying distinct habitat use patterns were differentially exposed to fishing capture. Finally, considering the unprecedentedly widespread and severe red tide and associated fish kill in the San Francisco Estuary in August 2022, we asked (4) during our study period spanning water years 2011-2017, what proportion of tagged white sturgeon tended to inhabit regions affected by the 2022 red tide event in late summer. Our results have implications for improved management of white sturgeon in the San Francisco Estuary and Sacramento River ecosystem.

## Methods

### Study System

The Sacramento River originates in the Klamath Mountains near Mount Shasta and flows through the northern portion of California’s Central Valley before joining the San Joaquin River near Antioch, CA. This confluence forms the Sacramento-San Joaquin River Delta, which then flows through Suisun and San Pablo Bays before reaching the San Francisco Bay (Figure 1). In addition to flowing through intensely urbanized and agricultural areas, the system has undergone substantial human modification, including draining of wetlands, levee construction, channel dredging, and water diversion for agricultural and municipal uses (Andrews et al. 2017). Further, the Delta is the heart of California’s massive water conveyance system, used to redistribute water across the landscape, in part, to irrigate a highly productive agricultural region. Inhabited by many declining populations of native fishes, including white sturgeon, green sturgeon, Delta smelt, longfin smelt and Chinook salmon (Moyle et al. 2011), this system has also been heavily instrumented with acoustic telemetry receivers to facilitate study and management of multiple species (Singer et al. 2013; Michel et al. 2015; Colborne et al. 2022). Across the range of white sturgeon, many populations are in decline due to overfishing and habitat degradation (Hildebrand et al. 2016), and the extent of state and federal protections varies among regional subpopulations (Schreier et al. 2013; Hildebrand et al. 2016). The Sacramento-San Joaquin population is not currently federally protected, but there is evidence that this population is in decline (Hildebrand et al. 2016; Blackburn et al. 2019).

**Figure 1.**
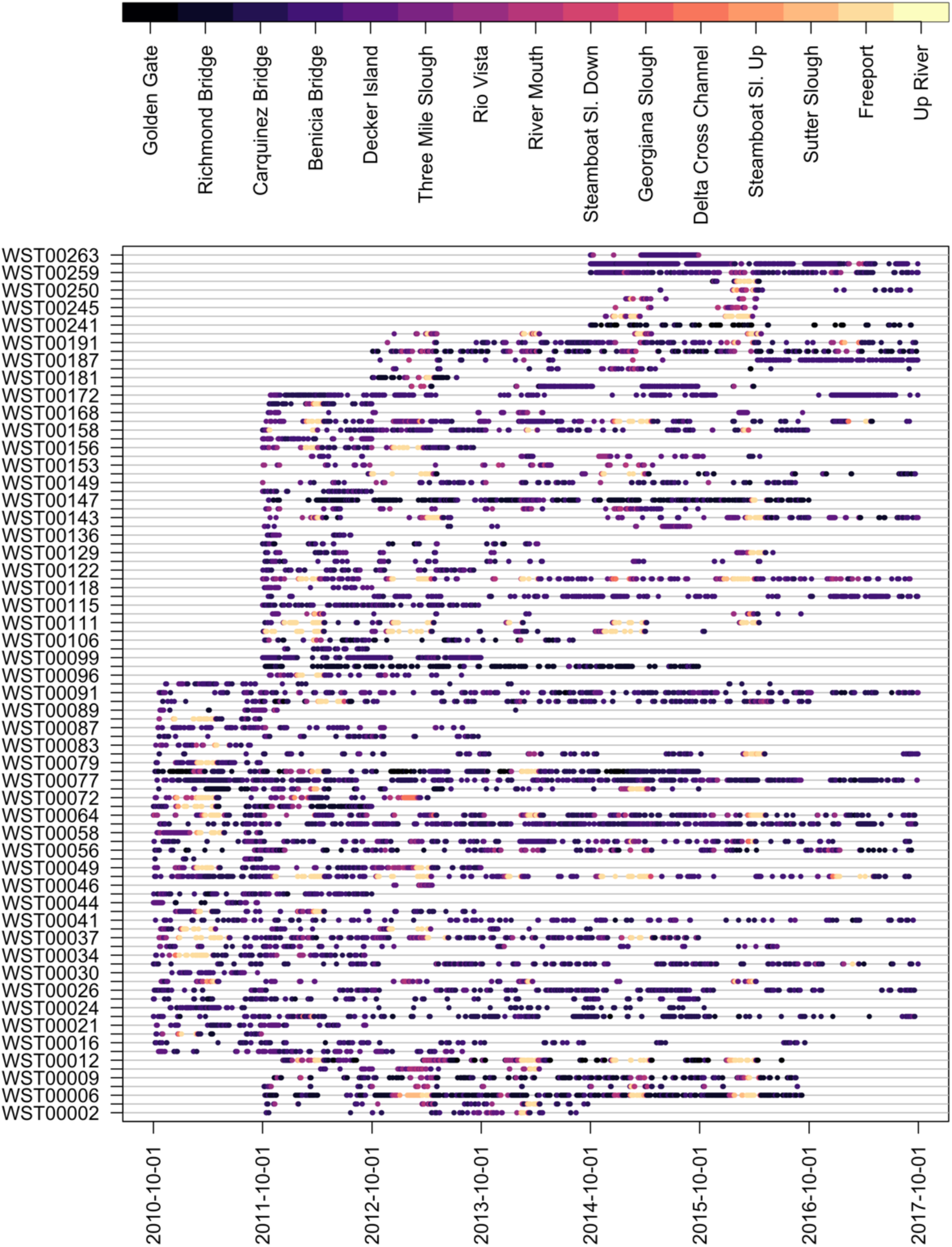
Detection histories of studied white sturgeon. Detection locations are ordered from bay mouth (Golden Gate Bridge; dark purple) to upstream portions of the Sacramento River (Up River; light yellow).

**Figure 2.**
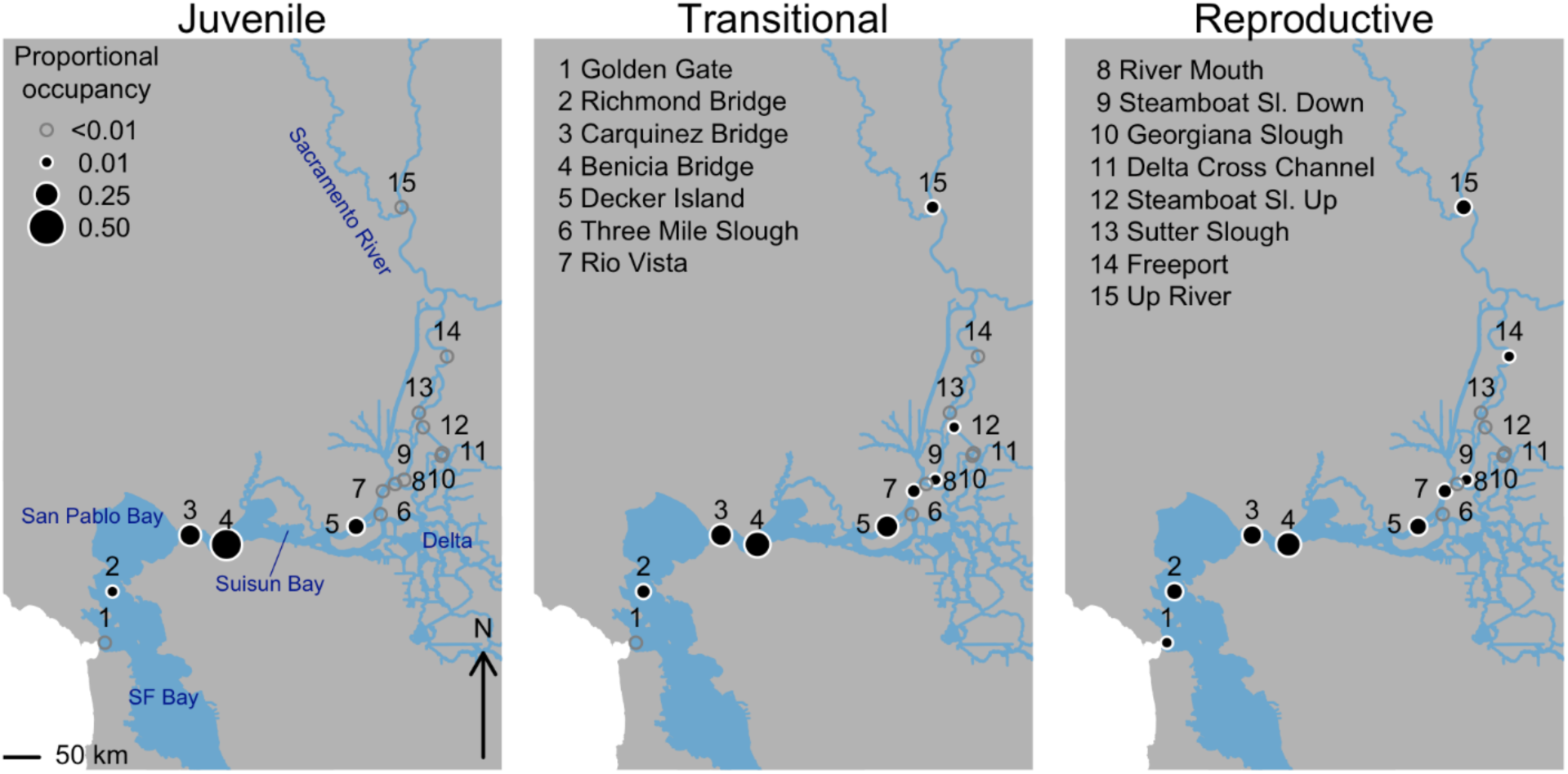
White sturgeon mean proportional occupancy by detection location for juvenile, transitional, and reproductive-age fish.

### White sturgeon telemetry data

We used data from three acoustic telemetry studies of white sturgeon contained in the Pacific Aquatic Telemetry Hub (PATH) database (<https://fishdb.wfcb.ucdavis.edu>). Capture methods varied depending on original study protocols, while tagging methods remained nearly identical across the three studies. Methods are described in detail in Miller et al. (2020), Johnston et al. (2020) and Thomas et al. (2013). Briefly, large juvenile and adult white sturgeon were captured with gill and trammel nets (Miller et al. 2020) in San Pablo Bay, Suisun Bay, and Grizzly Bay; rescued from the concrete channels behind the Tisdale and Fremont weirs, where they had been stranded by receding flood flows (Thomas et al. 2013); or captured in an eight-meter long, three-chambered fyke trap deployed in the toe drain of the Yolo Bypass (Johnston et al. 2020). All fish were tagged in the field upon capture and released nearby following the tagging procedure. Each fish was surgically implanted with an acoustic transmitter (V16, V13, or V9; Vemco, Ltd., Halifax NS) while inverted in a V-shaped cradle with continuous water flow across the gills. This tagging method has no significant impact on juvenile sturgeon survival, growth, or swimming performance (Miller et al. 2014).

During the 2011–2017 water years (WY; the California WY runs from October 1 to September 30) >300 acoustic receivers (VR2W-69 kHz, Innovasea Inc., Halifax NS, Canada) were deployed at 94 distinct sites through the collaboration of NOAA Fisheries, US Bureau of Reclamation, US Fish and Wildlife Service, ECORP Consulting, Inc, California Department of Water Resources, and US Geological Survey. In most cases, receivers were not deployed at the same location for the entire WY 2011–2017 period, so we focused on data from 15 sites in the Bay-Delta and Sacramento River system where at least one receiver was deployed on ≥90% of days from 1 October 2010 to 30 September 2017 (Figure 1). Receivers at two closely spaced sites (200 m) at the upstream intersection of the Sacramento River and Steamboat Slough were combined. All receivers deployed north of the confluence of the Feather and Sacramento rivers (ca. 38.76° latitude) were aggregated into one site we denoted as “up river.” There were also 8 detection locations on the San Joaquin and Mokelumne Rivers that met our criteria for near-complete deployment during the study period, but these were excluded from further analysis because detections of studied fish, while non-zero at these sites, were proportionally extremely uncommon. Excluding these sites had no substantive impact on our results.

There were 249 sturgeon detected during WY 2011-2017, and of these we included 106 in this analysis. Prior to selecting data for analysis, detection records were filtered to remove suspected false detections by omitting detections reflecting a single detection at a given site. In other words, detections were considered valid if a fish showed ≥2 consecutive detections at a given site. We discarded records from 4 individuals suspected to have perished when their detection records ended with a run of ≥200 detections/day at the same location for ≥30 days. These quantitative criteria were determined from visual inspection of individual fish detection records for records ending on long runs of repeated detections at the same location, without undetected periods of a day or longer suggestive of a fish moving away from and back toward a location. Because we could not be certain when a fish perished and because these cases were rare compared to the fairly large number of fish records available, we omitted all detections of these individuals from this analysis.

We selected from the remaining fish those with at least one complete water year in which the fish was detected on 30 different days. Water years were deemed complete if the fish was tagged no later than November 1 of the calendar year the water year began, and if the final detection of that fish was no earlier than September 1 of the calendar year in which the water year ended. There were 243 water years from 99 unique white sturgeon meeting these inclusion criteria. Selected fish were tagged in calendar years 2010 to 2014 in the studies described in Miller et al. (2020), Thomas et al. (2013) and Johnston et al. (2020). Although fish were tagged at different times of year (March through October) and there are seasonal patterns to white sturgeon movement (Miller et al. 2020), these influences were mitigated by analyzing only complete water years for a given fish. After filtering, we estimated detection probabilities (Supplementary Material S1).

### Occupancy analyses

We first examined how frequently white sturgeon were present at different sites and how this may have changed over time. We used telemetry detections to quantify, for each fish-water year combination, the number of fractional days that each fish was detected at each site. By fractional days, we mean that when fish were detected at *n*>1 site on the same day, the fish was considered to have spent 1/*n* days at each unique site, where *n* is the number of sites where the fish was detected on that day. This approach minimized the effect of fish being detected at multiple receivers in the same site, which could be biased by the number of receivers deployed at each site and changes therein over time. It also minimizes the effect of fish being detected many times in short succession by the same receiver array. More temporally explicit approaches, such as sequence analysis (Colborne et al. 2019), are poorly suited to this dataset because they require sampling over consistent time periods, whereas we used fish tagged in different years and having different lengths of detection record. To maximize the number of records studied in the present analysis, we used fish whose records spanned different time periods.

We quantified dissimilarity between all pairs of fish-water years in the proportion of days detected at each site and used surrogate- and permutation-based statistical tests to determine whether the observed differences in fish habitat use were greater than expected by chance. Dissimilarity was computed using Hellinger distance (Hellinger 1909). The Hellinger distance (*H*) between two individuals is defined as 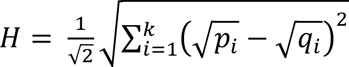, where here *p_i_* and *q_i_* are, respectively, the proportion of detections of fish *p* and *q* which occurred at location *i* in a given water year. Note that *p* and *q* may be the same fish in a different water year, different fish in the same water year, or different fish in different water years.

We conducted a surrogate-based statistical test to determine if the occupancy patterns were more dissimilar than expected by chance under a null hypothesis that fish did not differ in their probabilities of being detected at a given site, either across individuals or across water years. We constructed the null model by taking the mean proportional occupancy at each site over all fish-water years and using the mean occupancy profile to simulate a surrogate dataset that preserved the number of fish studied and the number of days each fish was detected in each water year. A surrogate occupancy profile for a given fish in a given water year was generated by randomly assigning a detection to a fish in a location and in a water year, based on the mean proportional occupancy profile constructed previously. Total numbers of detections of each fish in each water year were kept fixed at the values of the data. This process was repeated to create 1000 complete surrogate datasets. For the empirical dataset and each surrogate dataset, we calculated the total Hellinger dissimilarity among all pairs of two fish-water years and considered divergence in occupancy patterns to statistically differ from the null model of no divergence if the total Hellinger dissimilarity among all pairs of two fish-water years was greater than the 0.95 quantile of that statistic for the surrogates.

To test whether fish age class, fish sex, and water year flow conditions statistically explain divergence in occupancy behavior across fish and water years, we used permutational analysis of variance (PERMANOVA; McArdle & Anderson, 2001). PERMANOVA is used to partition dissimilarity matrices among sources of variation in a linear modeling framework and is commonly used in ecology for analyzing differences in community composition (e.g., Hart, Turcotte, & Levine, 2019; Hanna, Raudsepp-Hearne, & Bennett, 2020). We tested whether each predictor explained residual variation in dissimilarity in occupancy behavior while accounting for tagging region and tagging year. Tagging region and tagging year were included to account for studies making different choices about where to tag fish and whether to collect sex data over different years. Only one focal variable (age class, sex, water year conditions) was tested at a time to avoid partitioning the data into groups with unacceptably low sample sizes.

For the PERMANOVA analyses, we considered three age classes of fish: juvenile, transitional, and reproductive adult. Fish age at tagging was estimated using growth curves from Blackburn et al. (2019) and extrapolated forward in time to assign an estimated age to each fish in each water year. Juvenile fish were those <10 years of age based on the Blackburn et al. (2019) curve, transitional fish were ages 10-15, and reproductive adults were ≥ 16 years of age. Chapman (1989) indicated that white sturgeon begin sexually maturing at 10 years of age, with ≈80% of fish sexually mature by age 16. Age class bounds were also informed by the harvest slot limit (40-60 in, or 102-152 cm) on the white sturgeon recreational fishery that was active during the study period; transitional fish were of harvestable size. We observed 21 water years from 15 unique juvenile fish, 110 water years from 58 unique transitional fish, and 112 water years from 48 unique reproductive adult fish.

Fish sex was considered as male (18 unique fish over 41 fish-water year combinations), female (13 fish, 28 fish-water years), or unknown (68 fish, 174 fish-water years) depending on whether the source study sexed fish. We used water year type designations from the California Department of Water Resources, which categorizes each water year on a 5-point scale: Critically Dry; Dry; Below Normal; Above Normal; Wet. Tagging region was considered either “Bay” (Grizzly Bay, San Pablo Bay, Suisun Bay) or “Sacramento River” (Fremont Weir, Tisdale Bypass, Yolo Bypass). Tagging locations were aggregated to region because of low sample sizes for individual sites.

To interpret how these variables influence habitat occupancy, we categorized fish-water year occupancy records based on the most statistically important variables (i.e., those explaining the most variance in occupancy, which were also those with the lowest *p*-values) and took the mean proportional occupancy by site over groups of those variables.

### Exposure to fishing

We next evaluated whether any groups, based on characteristics that statistically explained differences in habitat use, experienced differential exposure to fishing capture. Fishing included both harvested and released individuals; note that, at time of writing, harvest of white sturgeon is currently illegal but there was a legal slot-based fishery during the study period. To quantify spatial variation in recreational fishing capture, we analyzed white sturgeon angler catch report card data collected annually by the California Department of Fish and Wildlife (CDFW). All fishers catching white sturgeon were required to report the date of their catch, whether it was harvested or released, the length of kept fish, and in which of 29 “fishing regions” the catch occurred. We used these data for 2015 through 2020 to estimate the geographic distribution of catch-and-release and harvest-oriented fishing capture for white sturgeon. Over this period, reporting practices, number of licenses issued, and report card return rates were largely consistent. Although there is uncertainty in fishing capture due to incomplete reporting and poaching, we take these as the best available index of relative differences in fishing capture across habitat zones of interest. We calculated a fishing capture risk index for each fish (*C_i_*) by apportioning region-specific fishing capture to individuals based on their proportional occupancy,

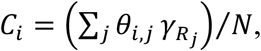

where *θ_i, j_* is the telemetry-determined proportional occupancy of fish *i* at detection location *j* in fishing region *R*, γ*_Rj_* is the estimated fishing capture in region *R* where detection location *j* is located, and *N* is the total number of white sturgeon captured. To calculate γ*_Rj_*, we averaged the reported number of captured sturgeon per region across 2015 to 2020. During 2015-2020, response rates were consistent (29.8% to 33.5%), and we did not attempt to further correct for incomplete reporting because reporting biases are likely but unquantifiable from currently available data. Despite uncertainties over the number of white sturgeon caught, we assert that these data provide valid first-order approximations of the spatial distribution of white sturgeon catch, which was our main objective. We then aggregated by age class to evaluate whether fishing capture risk tended to vary among age classes. We present results considering all caught (harvested + released) fish. Patterns in *C_i_* were similar when considering only harvested fish; however, since some age/size classes are not exposed to harvest, and because the sample of report cards was larger and hence less prone to sampling error when considering both harvested and released fish, we focus on results considering both.

### Potential red tide exposure

To estimate the proportion of the large juvenile and adult white sturgeon population exposed to the late-summer 2022 red tide, we used all white sturgeon detections from fish tagged in the selected studies during the 2010-2014 water years (including from fish not meeting selection criteria for the above analyses). The 2010-2014 water years were selected for this analysis because it was the period of greatest acoustic receiver coverage in areas affected by the 2022 HAB. We summarized these detection records to quantify the proportion of unique tagged fish detected in two focal sub-regions affected by the red tide (San Francisco Bay and San Pablo Bay) in 5-day-of-year intervals from January through December. By quantifying the proportion of unique tagged fish across coarse spatial and temporal scales, we minimize problems with multiple detections and potentially erroneous detections. A limitation of our approach, common in telemetry studies overall, is that receiver density and arrangement is inconsistent between and within regions: a fish may be more likely to be detected in an area of comparably high receiver density. However, many receiver arrays span bridges (e.g., Golden Gate, Bay Bridge, Benicia Bridge) so as to detect fish passing them at high rates. We estimated detection probabilities for our study, finding them to be high, 0.97 to 0.99, depending on analysis assumptions (Supplementary Material S1). Results are disaggregated by age class to examine whether different size/age groups have differing potential exposure risks based on their historical occupancy patterns.

All analyses were conducted in R version 4.0.3 (R Core Team 2021) using the ‘vegan’ (Oksanen et al. 2022), ‘abdiv’ (Bittinger 2020) and ‘stats’ (R Core Team 2021) packages.

## Results

### Habitat occupancy

Visual inspection of detection histories suggested that white sturgeon habitat use differs among individuals and water years (Figure 1). For example, certain fish (e.g., WST00006) migrated toward upstream spawning areas in some but not all water years, whereas others (e.g., WST00077) were never detected in upstream areas despite being observed over a similar time period. A permutation-based significance test corroborated that dissimilarity in habitat use of white sturgeon across individuals and water years was greater than was expected to arise by chance alone; the empirical total dissimilarity in habitat use profiles was greater than all 1000 surrogates (*p* < 0.001).

Fish age class (*df* = 2, 236; *F* = 2.51; *p* < 0.0001), but not fish sex (*df* = 2, 236; *F* = 1.63; *p* = 0.08) or water year flow conditions (*df* = 3, 242; *F* = 0.75; *p* = 0.73) statistically explained differences in habitat use behavior. Large juvenile fish were detected most frequently in the lowest portions of the Bay-Delta, with a modal detection location in Carquinez Strait (detection location 4, Benicia Bridge), and were detected anywhere upriver of Rio Vista (detection location 7) on < 1% of detections (Figure 1a). Transitional-stage fish were detected at a wider range of locations than juveniles, including in areas upriver of the confluence of the Sacramento and Feather rivers (detection location 15; Figure 1b). Reproductive adult were more likely than other size classes to be detected in areas upriver of the confluence of the Sacramento and Feather rivers (Figure 1c), which includes known spawning locations (Schaffter 1997). Reproductive adult white sturgeon were also observed at the mouth of San Francisco Bay (detection location 1, Golden Gate Bridge) with greater frequency than other size classes, suggesting that older/larger fish more commonly make trips into the Pacific Ocean. White sturgeon were detected at the Golden Gate Bridge in 23 fish-water year combinations; 15 of these were reproductive adults, 8 were transitional, and none were juveniles. There was also no significant effect of sex on differences in habitat use behavior when excluding individuals of unknown sex (*df* = 1, 65; *F* = 1.12; *p* = 0.33).

Release region and tagging year had statistically significant effects in all models, showing the importance of controlling for these factors while testing for influence of key variables. Yet, despite the statistically significant effects of size class, release region, and tagging year, these statistical models explained 9% - 22% of differences in total habitat use.

### Exposure to fishing

Fish in different age classes experienced differing levels of exposure to fishing capture and harvest. By merging biotelemetry-based habitat use information with angler report card catch data, we showed that transitional-age fish, which also were estimated to be of harvestable size during the study period, were less likely than other size classes to be detected in areas where most sturgeon are caught (Figure 3a). By far, most white sturgeon were captured by anglers in Suisun Bay and the lower Sacramento-San Joaquin Delta (Figure 4). This result does not appear to be driven by biases wherein our data include only fish surviving a full water year at harvestable size. There were no differences among age classes in the probability of including in our analyses consecutive water years of data for the same fish (binomial GLM; df = 2, 240, *p* = 0.75); and the frequency of including in our analyses at least one water year of data for the same fish both below the slot limit and at harvestable size (7 out of 15 fish) was indistinguishable from the overall probability of including in our analyses the same fish in successive water years (≈0.5).

**Figure 3.**
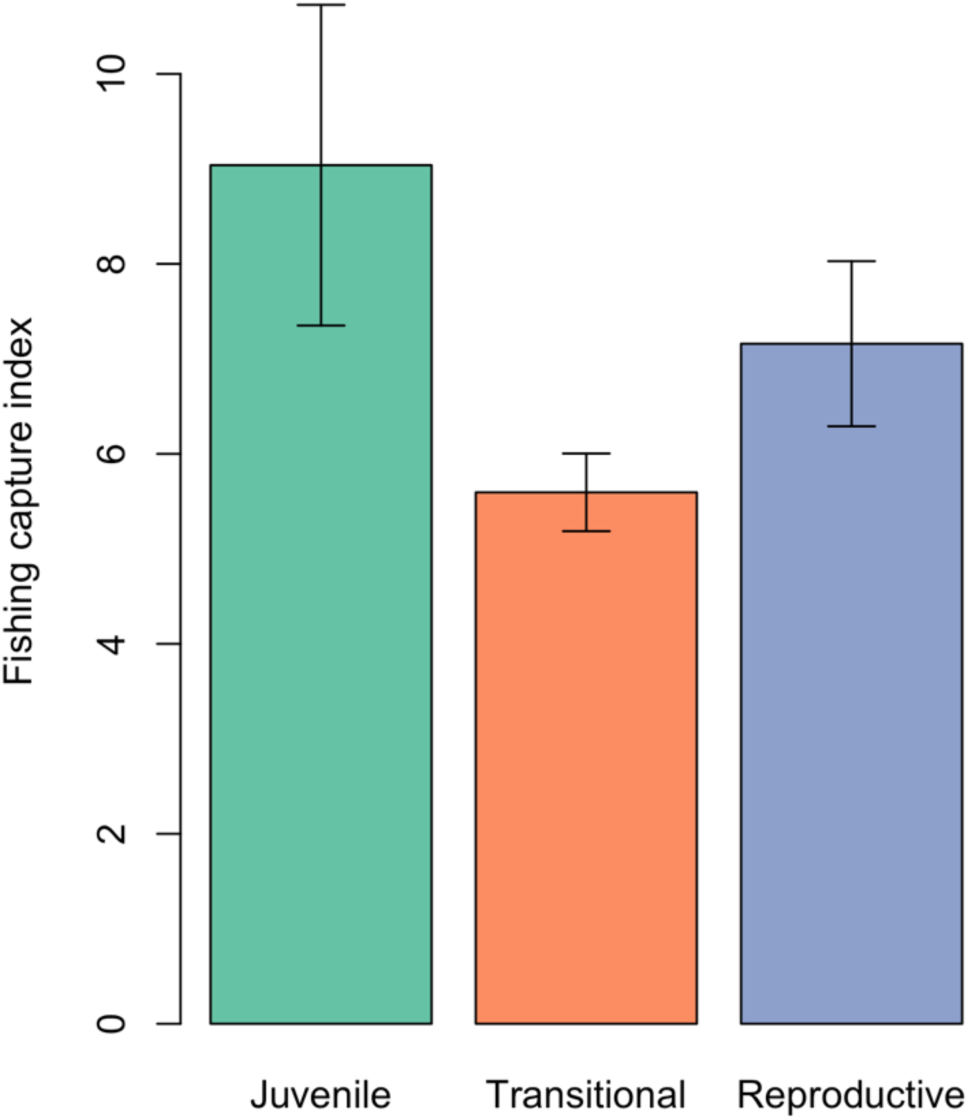
Index of exposure to fishing capture by age class based on self-reported data on the location of caught white sturgeon from 2015 to 2020. Error bars indicate one standard error.

**Figure 4.**
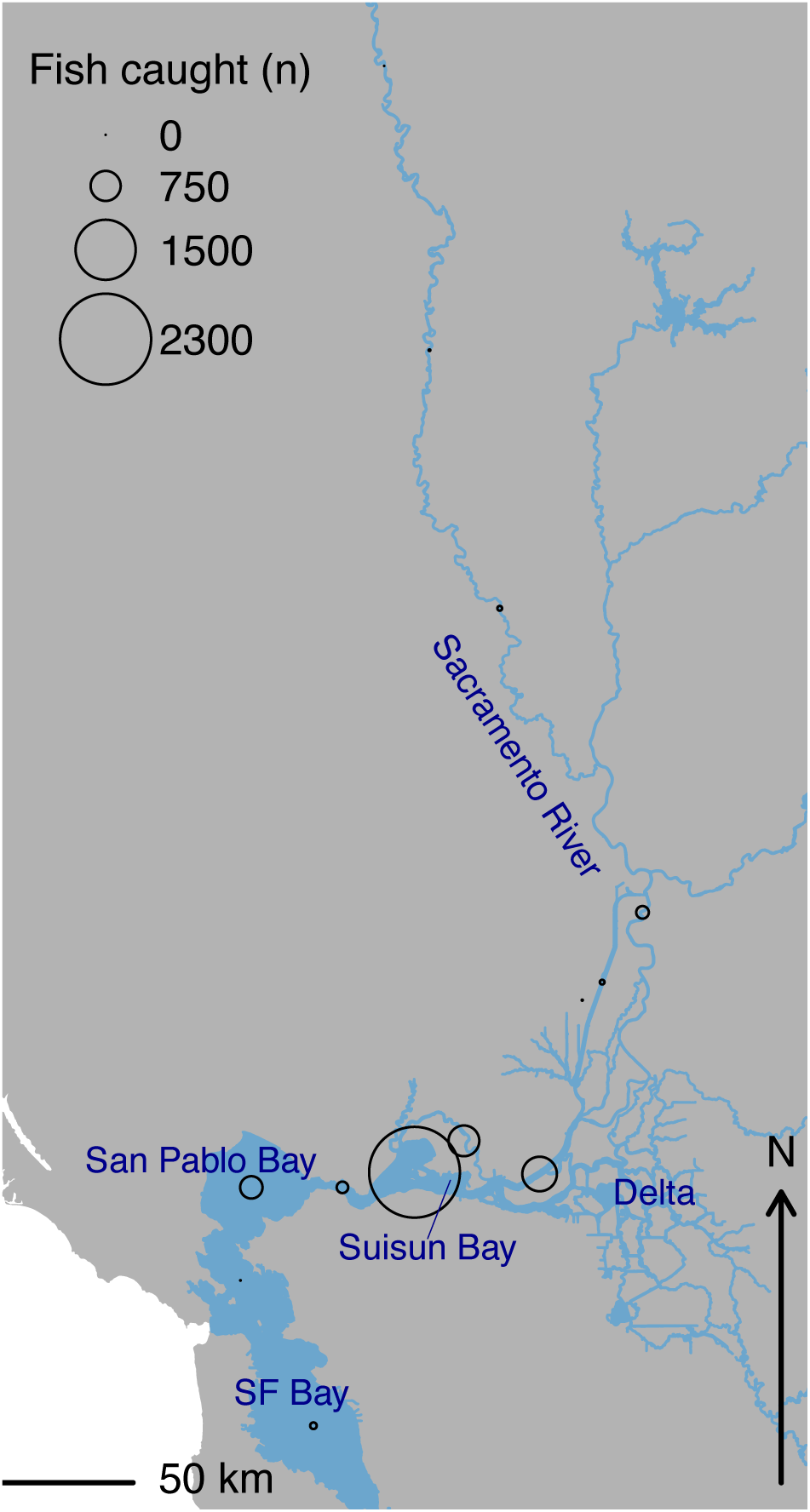
Estimated annual mean number of white sturgeon caught (harvested + released) by reporting region. Data are from 2015 to 2020. Regions are represented by points at their approximate centroids or midpoints due to their differing geometries (e.g., bays versus river segments).

### Potential red tide exposure

Based on historical patterns of habitat use, a substantial fraction of white sturgeon may have been present in portions of the study area most severely affected by the late summer 2022 HAB at the time of its occurrence (Figure 5). This pattern depended on age class such that transitional and reproducing sturgeon were more likely to have been detected in the affected regions (Figure 5). Particularly for reproductive adult white sturgeon, around 45% were detected in San Pablo Bay and 10% in San Francisco Bay. While juvenile fish may have experienced little exposure to the red tide due to their limited use of these habitat areas during the affected season, potential exposure increased with size/age class.

**Figure 5.**
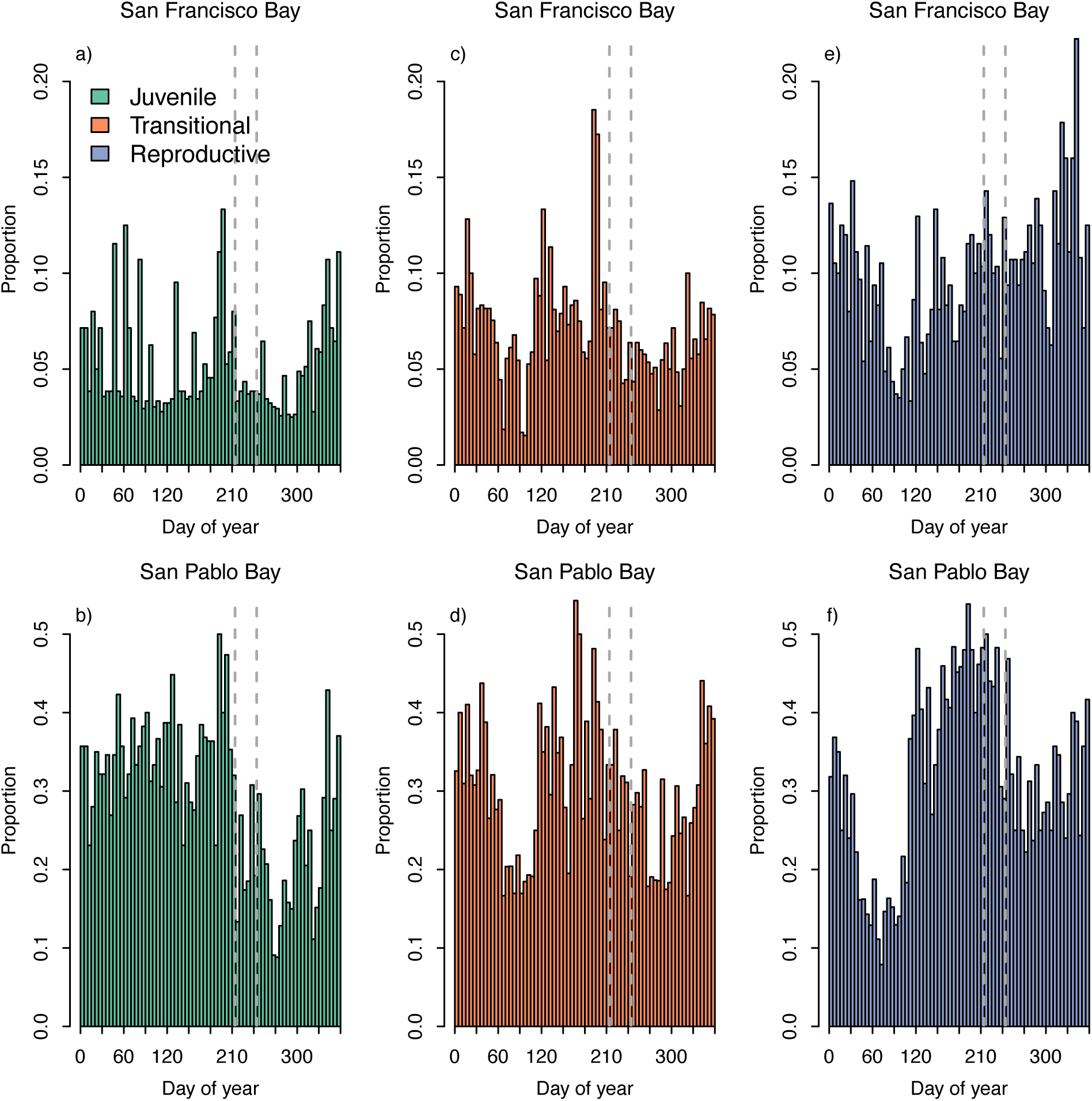
Proportion of fish at three size classes detected in two areas affected by harmful algal blooms (HABs) in summer 2022, San Francisco Bay and San Pablo Bay. Dashed vertical grey lines indicate the general period when the HAB occurred.

## Discussion

We conducted a synthesis of available acoustic telemetry data on white sturgeon to differentiate patterns of habitat occupancy across size classes in the San Francisco Bay, Sacramento-San Joaquin Delta, and Sacramento River system. Large juvenile white sturgeon (7-9 years est. age) concentrated occupancy in the upper bays and lowermost portions of the delta. Larger white sturgeon in the transitional and reproductive adult classes were detected increasingly frequently at furthest upriver sites that included known spawning areas, consistent with these age classes engaging in spawning migrations. The frequency of detections at the mouth of the San Francisco Estuary, possibly signaling movements into the Pacific Ocean, also increased with age class. We combined these data with other relevant data on fishing and a severe harmful algal bloom to improve understanding of sturgeon exposure to potential population threats. These analyses have implications for conservation management of white sturgeon and demonstrate an approach for synthesis science that leverages available long-term biotelemetry, fishery, and water datasets.

Our finding that white sturgeon at a transitional age class that also corresponded with the fishery slot limit were detected less frequently in areas where most white sturgeon are caught, as compared to smaller/younger (juvenile) or larger/older (reproductive adult), has multiple potential explanations. One logical but likely spurious possibility is that sturgeon are simply harvested once they reach the slot limit, giving them fewer opportunities to be detected in areas of high fishing pressure than out-of-slot fish, but this is unlikely considering how we selected near-complete fish-water year combinations for analysis and did not detect substantial biases toward selecting survivors for analysis. Alternatively, anglers may simply fail to direct their efforts toward areas where harvestable white sturgeon are most prevalent, for example if other factors determine preferences for where to fish (Dabrowksa et al. 2017; Hunt et al. 2019). Another possibility is behavioral avoidance. Although, to our knowledge, behavioral avoidance of anglers has not been studied in white sturgeon, other sturgeon species exhibit predator-avoidance behaviors and show signs of experiential learning even at age-0 (Hintz et al. 2013). Other fishes also can use prior experience and adjust behavior to avoid capture by anglers (Louison et al. 2019; Koeck et al. 2020). For behavioral avoidance alone to explain this result, not only must white sturgeon recognize and avoid areas of higher angler pressure, but this response must also be strongest in fish of harvestable size. While the former seems plausible, a mechanism making the angler-avoidance response strongest in harvestable white sturgeon seems unlikely. The hypothesis that habitat use of white sturgeon reflects behavioral responses to angling could be tested more explicitly in future research. Although harvest of white sturgeon is presently illegal in California, fishery regulations are dynamic and understanding the spatial distribution of angler pressure and how it intersects with sturgeon habitat use across life stages can inform future regulation of the white sturgeon fishery.

The late-summer 2022 toxic red tide and associated fish kill likely had a profound impact on the white sturgeon population, as evidenced by >800 sturgeon (green, white, and unidentified) carcasses recovered following the event. Our analyses, based on historical data several years preceding the red tide, suggested that substantial proportions of the white sturgeon population were likely residing in areas of the system affected by the red tide (i.e., San Francisco Bay and San Pablo Bay) at the time of year when the event occurred (Figure 5). This was particularly true of reproductive adult fish. While total mortality and mortality rates are uncertain, our findings suggest that >50% of reproductive-age fish could have been exposed to deadly red tide conditions based on their historical habitat occupancy patterns. Due to a combination of water body eutrophication and climate change making conditions supportive of harmful algal blooms like red tide more common, there is concern for increases in the frequency and severity of such events in the San Francisco Estuary (Kudela et al. 2023) and many other water bodies (Carstensen et al. 2007; Fey et al. 2015; Gobler 2020; Hou et al. 2022).

Our result that habitat use patterns change with sturgeon age complements another recent study of white sturgeon in the San Francisco Estuary and Sacramento-San Joaquin river system, but using different methods. Sellheim et al. (2022) used fin ray geochemistry to infer age-specific habitat use across a salinity gradient from open ocean to freshwater, with the average salinity inhabited by a fish tending to increase over ages 0 to 10. Fish in this age range are mostly juveniles, and in our study these tended to be found lowest in the system, in areas tending toward higher salinity, compared with larger fish. Note that the youngest fish in our dataset was estimated at 7 years old, which Sellheim et al. (2022) indicate would tend to be using higher-salinity habitats. The discrepancy in ages between these studies also highlights that telemetry data on young white sturgeon are limited, resulting in uncertainty over their habitat use and movements. While studies of other white sturgeon populations have generally found differences among fish in habitat use and movement (Parsley et al. 2008; Nelson and McAdam 2012; Schreier et al. 2013; McLean et al. 2020), direct comparison is challenging because most other studies have been focused on more local spatial scales or have not examined variation across life stages.

Despite our results showing habitat use differences by age class, substantial variability in habitat use behavior was unexplained and could be associated with other factors. Related species exhibit habitat use differentiation into groups of individuals with characteristically different occupancy patterns that are not clearly associated with life stage or sex. Migratory populations of native lake sturgeon (*Acipenser fulvescens*) also exhibit spatiotemporally segregated “contingents” with distinct patterns of habitat use in the Laurentian Great Lakes region, including resident and migrant individuals as well as distinct migration patterns (Kessel et al. 2018; Colborne et al. 2019). For lake sturgeon, habitat use patterns were generally consistent over time, suggesting that sturgeon habitat use groups reflect individuals’ intrinsic preferences or specialization, possibly influenced by life history factors. Groups of shortnose sturgeon (*Acipenser brevirostrum*) inhabiting coastal waters and river systems of Maine also exhibit divergent patterns of movement and habitat use (Dionne et al. 2013; Altenritter et al. 2018), and in the same Sacramento River system green sturgeon (*Acipenser medirostris*) exhibit distinctly bimodal early/late downriver migration patterns (Colborne et al. 2022). Sellheim et al. (2022) found evidence for groups of white sturgeon from the same population as this study exhibiting distinct patterns of habitat use with age. Intraspecific variation in habitat occupancy and migration behavior could be a common and linking feature of sturgeon ecology more generally (Miller et al. 2020). The potential benefits of divergent behaviors for bet-hedging against local catastrophe may be important stabilize sturgeon populations, particularly considering that other life history characteristics that facilitate recovery from disturbance, such as rapid population growth, are not present in sturgeon.

Our findings have important implications for white sturgeon monitoring and management in the San Francisco Estuary and Sacramento River system. Approaches to monitoring white sturgeon population health in the San Francisco Estuary have recently shifted, replacing a single-location trammel-net survey with a spatially balanced, fishery-independent, mark-recapture study and spawner surveys. The new mark-recapture study samples at many locations following a stratified random design. Our study supports the importance of a structured, multi-location survey for limiting potential biases that could arise from sampling in a single location, as well as for strengthening understanding of spatiotemporal variation in white sturgeon densities and individual movements. We found evidence that a considerable fraction of the white sturgeon population likely could have been exposed to red tide conditions in August 2022. We were unable to address in greater detail how the red tide event affected sturgeon behavior or mortality rates due to limits in data availability, including receiver deployments, though it’s clear that population impacts were substantial. In anticipation of future HABs, expanding receiver deployments in the San Francisco Estuary could produce new, more spatiotemporally detailed data on white sturgeon habitat use and responses to HABs. By leveraging an existing, broadscale, multi-agency acoustic telemetry array (Cordoleani et al. 2021), we can continue to develop new insights, with relatively low additional costs, by continuing to equip white sturgeon with long-lived acoustic transmitters and maintaining an array of acoustic receivers. Continued long-term telemetry studies and monitoring, such as presented here, can provide critical evidence and support toward improving management practices.

In the San Francisco Estuary and Sacramento River system, white sturgeon in different age classes exhibited variable patterns of habitat use. Differentiated patterns of habitat use and movement within populations appear to be a common feature of sturgeon biology (Dionne et al. 2013; Kessel et al. 2018; Altenritter et al. 2018; Colborne et al. 2019, 2022; Sellheim et al. 2022). Further study is needed to resolve the biological underpinnings and implications for population health and management. Our findings suggest that different age classes of white sturgeon are exposed to different levels of risk from recreational fishing and harmful algal blooms, but the extent to which white sturgeon movement behaviors respond to these factors is not fully resolved. One further consideration for white sturgeon conservation is how habitat preferences intersect with boating traffic. Boat strikes are a documented cause of white sturgeon mortality in this region (Demetras et al. 2020), but research is needed to quantify the rate and spatial distribution of boat strikes of white sturgeon to better understand the impacts of boat strikes on the population and, if warranted, to inform possible regulation. This study demonstrates the value of synthesis research using long-term tracking studies, particularly for understanding the ecology of organisms such as sturgeon that are long-lived, mobile and use a diversity of habitats across their lifespans.

## Statements and Declarations

### Open science

Raw acoustic telemetry detection data used in this manuscript are publicly accessible on the PATH database (<https://fishdb.wfcb.ucdavis.edu>). Analysis code and supporting and derived datasets are available during review at <https://github.com/jonathan-walter/wstelemetrythreats-ms> and will be permanently archived on Zenodo and assigned a DOI upon manuscript acceptance; the Zenodo DOI will be provided in the published version.

## Acknowledgements

This analysis was supported by California Delta Science Programs grant 18204. This work was also supported by the University of California, Davis Agricultural Experiment Station (grant #2098-H to NAF and grant #2467-H To ALR). ALR was also supported by the Peter B. Moyle and California Trout Endowment for Coldwater Fish Conservation. Fish tagging was supported by funding and/or in-kind support from the U.S. Army Corps of Engineers and California Department of Fish and Wildlife. The receiver array is maintained through the collaboration of NOAA Fisheries, USBR, USFWS, ECORP, CDWR, and USGS. We also would like to thank the myriad individuals who tagged sturgeon and maintained receiver arrays, including the Biotelemetry Laboratory at UC Davis, and partners at CDFW, USACE, USBR, NOAA Fisheries, USFWS, CDWR, ECORP, and USGS, whose efforts provided the extensive datasets used for this analysis. Colby Hause facilitated angler report card data access from CDFW.

## Supporting Information for

### S1: Estimating detection probabilities

We estimated detection probabilities in a portion of our study area where receiver placements relative to the structure of the bay, delta, and river network provided unambiguous paths that should be followed by all fish. The area was between Richmond Bridge (location 2 in Figure 2), Carquinez Bridge (location 3 in Figure 2), and Benicia Bridge (location 4 in Figure 2). Each bridge features an array of receivers along its span, and there are no significant movement paths between locations 2 and 4 that do not include passing location 3. Due to the braided nature of the Sacramento-San Joaquin Delta and the presence of water conveyance infrastructure like the Yolo Bypass, other suitable sets of locations with unambiguous movement paths between them did not exist. Since we analyzed the data at the level of sites, not individual receivers, we conducted our detection probability analysis at the same level of aggregation.

After cleaning the data to remove false detections and selecting fish-water year combinations for analysis as described in the main text, we identified detection events using the ‘glatos’ R package (REF). For robustness, detection events were calculated in two ways: once with a duration limit of 60 minutes, so that when a fish was detected at the same location multiple times spanning >60 minutes it would be considered >1 detection event (i.e., consecutive detection events can be at the same location); and once with no duration limit, so that only changes in location would be considered a new detection event. We estimated the detection probability as the fraction of times that, after a fish was detected at Carquinez Bridge, its next detection event was at Richmond Bridge, Benicia Bridge, or (when allowing consecutive detection events at the same location) again at Carquinez Bridge. The estimated detection probability when allowing consecutive detection events at the same location was 0.988, and when only considering changes in location was 0.969. There is some uncertainty how accurate this estimate is for other parts of the study area. For example, the sites used to estimate detection probabilities feature a “curtain” style array with several receivers placed along the span of a bridge, whereas other sites have fewer receivers in different arrangements. Nonetheless, all sites have receiver arrays designed for accurate detections given the geometry of the waterbody at that site.

